# Native mass spectrometry can effectively predict PROTAC efficacy

**DOI:** 10.1101/851980

**Authors:** Rebecca Beveridge, Dirk Kessler, Klaus Rumpel, Peter Ettmayer, Anton Meinhart, Tim Clausen

**Affiliations:** Research Institute of Molecular Pathology (IMP), Vienna Biocenter (VBC), Campus-Vienna-Biocenter 1, 1030 Vienna, Austria; Discovery Research, Boehringer Ingelheim Regional Center Vienna GmbH & Co KG, 1120 Vienna, Austria

## Abstract

Protein degraders, also known as proteolysis targeting chimeras (PROTACs), are bifunctional small molecules that bring an E3 ubiquitin ligase and a protein of interest (POI) into proximity, thus promoting ubiquitination and degradation of the targeted POI [1–3]. Despite their great promise as next-generation pharmaceutical drugs, the development of new PROTACs is challenged by the complexity of the system, which involves binary and ternary interactions between components. Here, we demonstrate the strength of native mass spectrometry (nMS), a label-free technique, to provide novel insight into PROTAC-mediated protein interactions. We show that nMS can monitor the formation of ternary E3-PROTAC-POI complexes and detect various intermediate species in a single experiment. A unique benefit of the method is its ability to reveal preferentially formed E3-PROTAC-POI combinations in competition experiments with multiple substrate proteins, thereby positioning it as an ideal high-throughput screening strategy during the development of new PROTACs.

---

The development of PROTACs, small molecule “protein degraders” (**Fig. 1a**), is an emerging strategy in drug discovery, having major advantages over traditional small molecule inhibitors. PROTACs eliminate a target protein rather than inhibit it and function in a catalytic manner, requiring sub-stoichiometric amounts to achieve efficiency [4]. Moreover, PROTACs are applicable to a wider spectrum of proteins since degradation is not limited to a specific functional domain or active site [5]. To date, protein degraders have been developed against a variety of medically relevant proteins, such as the tumorigenic Androgen Receptor and Estrogen Receptor, as explored in first clinical trials [5–10]. In order to realise the full potential of protein degraders, methods have been developed to address the complex kinetics of multi-component PROTAC systems, which comprise various intermediate states [11–13]. Notably, the analysis of ternary interactions requires certain approximations to overcome the limitations of traditional biophysical techniques and always demands multiple experiments to estimate the basic kinetic parameters of the PROTAC system of interest.

**Figure 1.**
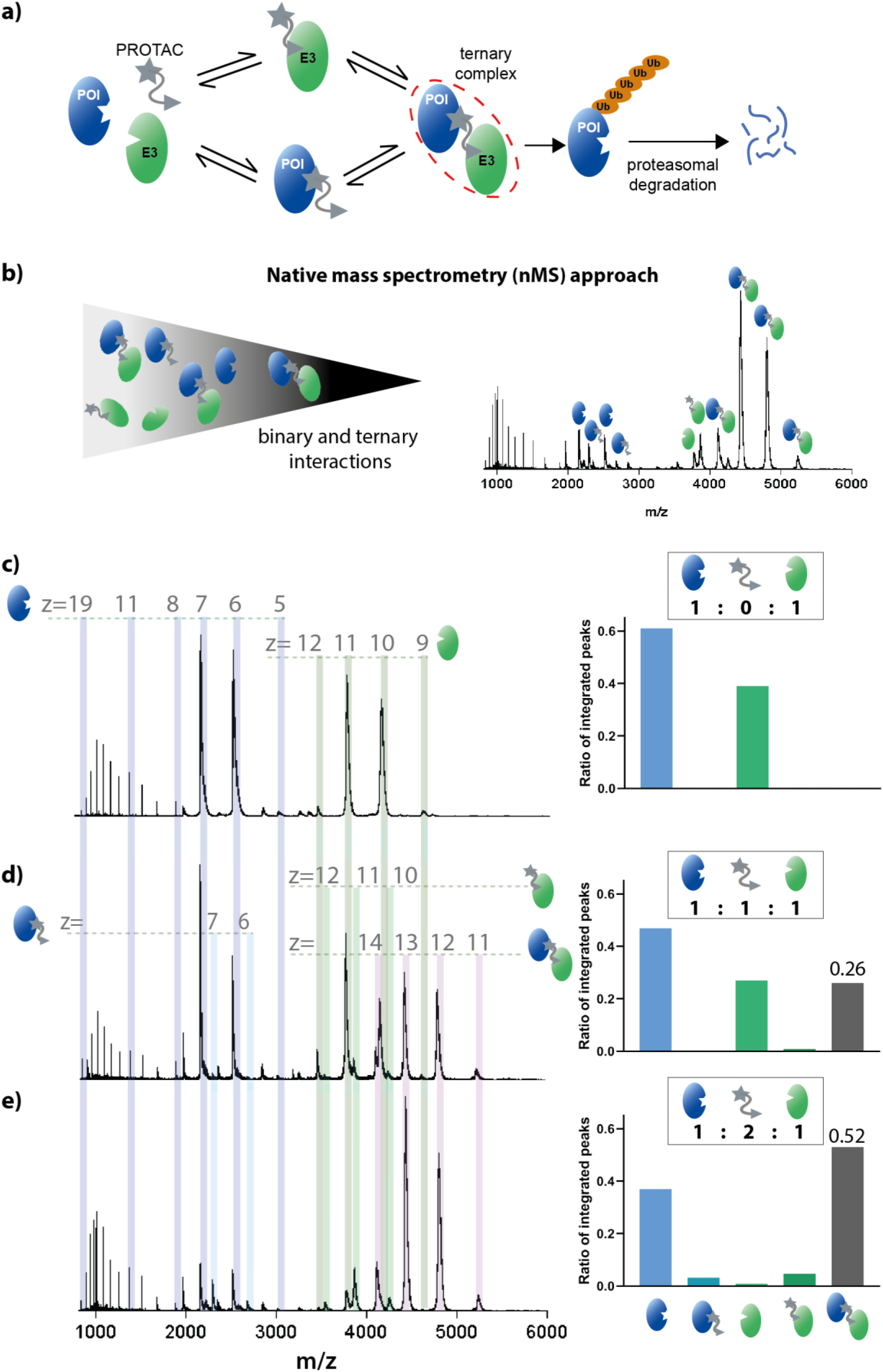
(a) Mechanism of targeted protein degradation by a proteolysis targeting chimera (PROTAC) molecule. The substrate protein is shown in blue and the E3 ligase in green. (b) Schematic of the native mass spectrometry (nMS) approach. (c-e) nESI-MS of Brd4^BD2^ (5 μM, 15 036 Da) and VCB (5 μM, 41 376 Da) sprayed from ammonium acetate (100 mM, pH 6.8) and 0.5% DMSO at AT1 (971 Da) concentrations of 0 μM (c), 5 μM (d) and 10 μM (e). Insets show estimated relative signal intensity of integrated peaks corresponding to *apo-*Brd4^BD2^, binary AT1-Brd4^BD2^ complex, *apo-*VCB, binary AT1-VCB complex and ternary complex Brd4^BD2^-AT1-VCB. Bar charts are representative of a single measurement, so no error bars are shown in this case. Expected and measured masses of each species are reported in **Table S1**. At an equimolar ratio, the signal intensity of Brd4^BD2^ is higher than that of VCB, likely due to higher ionisation efficiency as a result of its smaller mass and higher charge states.

With the use of nano-electrospray ionisation (nESI) [14], protein complexes can retain their native topology and stoichiometry during transfer from solution into the gas phase, making protein-protein and protein-ligand interactions amenable to MS analysis [15]. Key advantages of this “native MS” approach [16] include the label-free measurement of protein complexes and its capability to report on multiple binding stoichiometries present in dynamic protein mixtures, including molecular species populated to a low extent [15, 17–20]. For these reasons, we anticipated that nMS would be particularly applicable for the characterisation of PROTAC systems. It could complement quantitative biophysical methods such as isothermal titration calorimetry (ITC) and surface plasmon resonance (SPR) by analysing the E3, PROTAC and POI interplay in a single experiment (**Fig. 1b**). Here, we demonstrate that nMS can (1) report on the formation of E3-PROTAC-POI ternary complexes in a semi-quantitative manner, (2) delineate the binding specificity of a particular PROTAC molecule and (3) simultaneously measure PROTAC specificity to multiple substrate proteins in a single measurement. To this end, we used the two established PROTACs AT1 and MZ1, which target bromodomain-containing proteins for degradation via the Von-Hippel-Lindau (VHL) E3 ligase, as model compounds. Specificity, affinity and degradation behaviour of AT1 and MZ1 towards different bromodomains has been well characterised [12, 21–23], providing an excellent test system to benchmark nMS as an analytical tool in PROTAC research. Substrate proteins investigated include the first and second bromodomains of Brd4 (Brd4^BD1^ and Brd4^BD2^), and the second bromodomain of Brd3 (Brd3^BD2^). E3, PROTAC (P) and substrate (S) combinations that were characterised by MS are summarised in **Table 1**, together with results from previous ITC and SPR analyses.

**Table 1.** Comparison of native MS data on ternary complex formation with literature values.

|  |  | K<sub>D</sub> of VCB binding to [PROTAC + substrate] |  | Cooperativity | t<sub>1/2</sub> of ternary complex | fraction of ternary complex |
| --- | --- | --- | --- | --- | --- | --- |
|  |  | ITC [21] | SPR [12] | SPR [12] | SPR [12] | nMS |
| PROTAC | substrate |  |  |  |  |  |
| AT1 | Brd4<sup>BD1</sup> | 390 nM | 578 nM | 0.2 | <1s | 0.65 ± 0.1 |
| AT1 | Brd3<sup>BD2</sup> | 79 nM | 163 nM | 0.7 | 3s | 0.58 ± 0.07 |
| AT1 | Brd4<sup>BD2</sup> | 46 nM | 24 nM | 4.7 | 26s | 0.82 ± 0.06 |
| MZ1 | Brd4<sup>BD1</sup> | 28nM | 30nM | 0.9 | <1s | 0.80 ± 0.06 |
| MZ1 | Brd3<sup>BD2</sup> | 7nM | 8nM | 3.6 | 6s | 0.83 ± 0.05 |
| MZ1 | Brd4<sup>BD2</sup> | 4nM | 1nM | 22 | 130s | 0.92 ± 0.03 |

We first tested the capability of nMS to resolve dimeric (E3:P, E3:S, P:S) and trimeric (E3:P:S) complexes present in the reaction mixture. This initial analysis was focussed on Brd4^BD2^ (S) and its interaction with the VHL/elongin-B/elongin-C complex (VCB, E3), with and without AT1/MZ1 (P). As reference, native mass spectra of VCB and Brd4^BD2^ (5 μM) were recorded separately, sprayed from 100 mM ammonium acetate, and 100mM ammonium acetate containing 0.5% DMSO (**Fig. S1 and S2**), the latter condition used in all experiments monitoring complex formation. Expected and measured masses of each species are provided in **Table S1**. A native mass spectrum of a VCB and Brd4^BD2^ mixture (**Fig. 1c**) shows that no interaction occurs between the proteins in the absence of a PROTAC molecule. VCB presents in charge states [M+9H]^9+^ to [M+12H]^12+^, and Brd4^BD2^ presents in charge states [M+5H]^5+^ to [M+17H]^17+^, with most of the intensity in charge states [M+6H]^6+^ and [M+7H]^7+^. The inset shows the estimated fractional ratios of the integrated peaks corresponding to each species, calculated by summing the intensity of each charge state corresponding to a particular species, and normalised to the summed intensity of all annotated peaks in the spectrum. Upon the addition of 5 μM AT1 (1:1:1 ratio of E3:P:S), peaks are present at m/z ratios corresponding to that of the ternary Brd4^BD2^-AT1-VCB complex. Compared to the signal intensity of the ternary complex (0.25 of total intensity), signal corresponding to the binary VCB-AT1 species (E3:P) is in very low abundance (0.01), while the Brd4^BD2^-AT1 species (P:S) is not observed at all. When the AT1 concentration was increased to 10 μM (1:2:1 ratio of E3:P:S), the signal corresponding to the ternary complex becomes dominant (0.55), compared to VCB-AT1 (0.05) and Brd4^BD2^-AT1 complexes (0.03) and the *apo* VCB (0.02). Only the signal corresponding to *apo* Brd4^BD2^ remains relatively high (0.35). Interestingly, upon addition of AT1, the intensity of the [M+6H]^6+^ and [M+7H]^7+^ charge states become lower compared to the charge states [M+8H]^8+^ and above. These lower charge states correspond to a compact, folded conformation, with the higher charge states corresponding to an unfolded protein subpopulation [24, 25]. We can therefore infer that the folded conformation of Brd4^BD2^ is incorporated into the ternary complex, whilst the unfolded subpopulation remains isolated. Consistent with this, the Brd4^BD2^-AT1 binary complex is present only in [M+6H]^6+^ and [M+7H]^7+^, and no peaks are present corresponding to higher charge states. Distinguishing between folded and unfolded factions present in a substrate sample further highlights the potential of nMS in characterizing complex PROTAC reaction mixtures. To compare complex formation with a different PROTAC, equivalent measurements were carried out with MZ1 (**Fig. S3**). The overall distribution of binary and ternary complexes is similar to those formed with AT1, but in this case no binary complex between Brd4^BD2^ and MZ1 is observed, even at 10 μM MZ1 concentration. These data fit to the higher stability of ternary complex initiated by MZ1 relative to AT1, as determined by SPR measurements (**Table 1**). In sum, the initial MS measurements demonstrate the strength of nMS for the characterisation of protein complexes formed by PROTAC molecules, providing a semi-quantitative description of the binding equilibrium between E3, substrate and PROTAC. Moreover, the method reveals characteristic differences in reaction intermediates formed with different PROTACs, implying mechanistic differences in ternary complex formation.

To investigate whether nMS can report on the specificity of PROTACs for particular substrate proteins, we took advantage of the preference of AT1 to form ternary complexes with Brd4^BD2^ over other bromodomain-containing proteins [12, 21, 22]. Spectra of a VCB:AT mixture with different bromodomain substrates Brd4^BD2^, Brd3^BD2^ and Brd4^BD1^ respectively, are shown in **Fig. 2 a-c** (spectra of isolated Brd4^BD2^, Brd3^BD2^ and Brd4^BD1^ in **Fig. S2**, **Fig. S4** and **Fig. S5**). Owing to the different ionization efficiencies of the free substrates, we integrated the signal intensity of *apo-*VCB, the binary VCB-AT1 complex and the three ternary complexes. Comparing the relative amounts of ternary complexes reveals the preferential engagement of Brd4^BD2^ by VCB:AT1 (0.82), relative to Brd3^BD2^ (0.58) and Brd4^BD1^ (0.65). These data are consistent with previous ITC experiments, where VCB was mixed with saturated PROTAC-substrate complexes to estimate the K_d_ of ternary complex formation [21] (**Table 1**). Although nMS and ITC measurement predict slightly different preferences in binding Brd3^BD2^ and Brd4^BD1^, both methods highlight that the VCB:AT1 system most favourably forms a ternary complex with Brd4^BD2^. We next investigated the PROTAC MZ1, which binds all bromodomain substrates with higher affinity than AT1, but displaying less selectivity for Brd4^BD2^ (**Fig. S9**). In this case, the relative amounts of ternary complexes with Brd4^BD2^, Brd3^BD2^ and Brd4^BD1^ are 0.92, 0.83 and 0.80, respectively, pointing to a similar stability of the formed complexes (triplicate measurements shown **Fig. S10-12**). For MZ1 the K_d_ values from ITC are 4 nM, 7 nM and 28 nM for Brd4^BD2^, Brd3^BD2^ and Brd4^BD1^, respectively, fitting nicely to the nMS data (**Table 1**). Taken together, the nMS data demonstrate the pronounced selectivity of AT1 towards Brd4^BD2^, whereas MZ1 is a less selective PROTAC targeting bromodomains.

**Figure 2.**
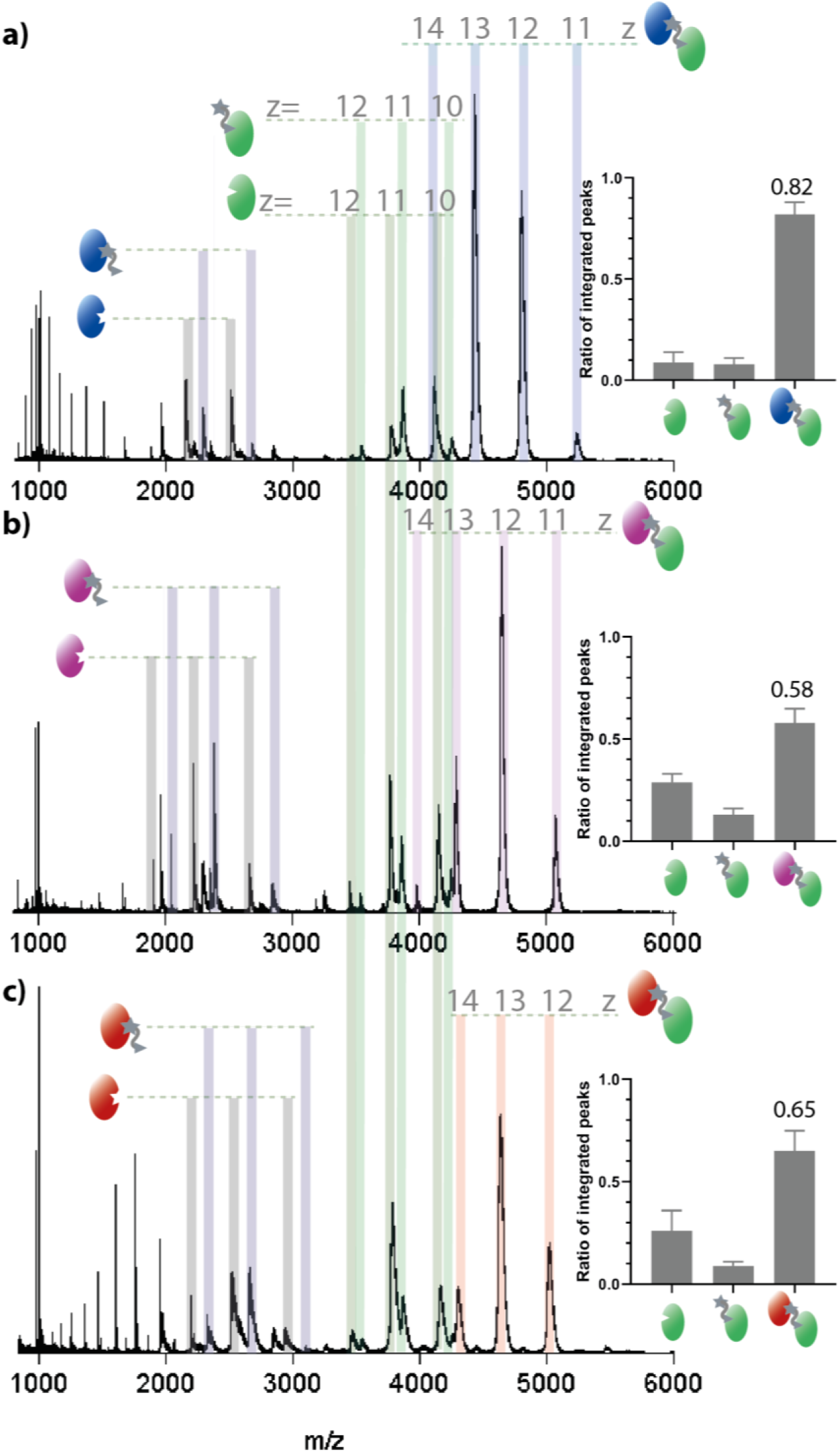
nESI-MS of VCB (5 μM), AT1 (10 μM) and Brd4^BD2^ (5 μM, a), Brd3^BD2^ (5 μM, b) or Brd4^BD1^ (5 μM, c). Proteins are sprayed from a starting solution of ammonium acetate (100 mM, pH 6.8) and 0.5% DMSO. Inset; estimated relative intensity of summed peaks corresponding to *apo-* VCB, binary AT1-VCB complex and ternary complex substrate-AT1-VCB. Samples were analysed in triplicate (**see Fig. S6-8**) and the error bars represent standard deviation of the relative peak intensity. Fractional intensity of signal corresponding to the ternary complex is shown, and values for other species can be found in **Table 1.**

An additional question regarding the mechanism of PROTACs is whether they display cooperative behaviour. To address this point, we measured E3:AT1 and S:AT1 mixtures and compared the binary complex formation to ternary complex formation, using a cooperative (Brd4^BD2^) and a non-cooperative (Brd4^BD1^) substrate (**Fig. S13**). Binary complex formation of VCB:AT1 is low, below 0.2, and the binary complex formation of Brd4^BD2^:AT1 and Brd4^BD1^:AT1 is roughly 0.5 in both cases. When the three components are mixed together, however, the ternary complex is formed to a much higher extent with Brd4^BD2^ (0.82) than Brd4^BD1^ (0.65), hinting at cooperativity of this PROTAC system, as proposed by [21]. According to these data, nMS analysis allows, in addition to determining the most favoured ternary complexes, to distinguish differences in cooperativity between PROTAC systems.

Finally, in order to take full advantage of the benefits of nMS over other biophysical methods, we applied our approach to a complex reaction mixture containing an E3, a PROTAC and multiple substrates. Since nMS was able to distinguish PROTAC specificity in separate experiments, we were curious to what extent PROTACs would recruit the bromodomains in a competition experiment that mimics the in vivo situation more closely. Ternary complex formation was measured using equimolar amounts of Brd4^BD2^, Brd3^BD2^ and Brd4^BD1^ and the PROTAC MZ1 that seemingly promotes ternary complex formation in a rather unselective manner (**Fig. S9**). Initially, an overall substrate concentration (S1+S2+S3) equimolar to that of VCB was used, thus avoiding competitive binding. In this case, the relative signal intensity of ternary complex incorporating Brd4^BD2^ is the highest, with that incorporating Brd3^BD2^ at a slightly lower intensity and that incorporating Brd4^BD1^ at an even lower intensity (**Fig. 3a**). When the substrate concentration is increased threefold, thereby increasing the competition for binding, the signal intensity of the Brd4^BD2^-containing ternary complex is more than three times higher than complexes containing Brd3^BD2^ and Brd4^BD1^ (**Fig. 3b**), clearly outcompeting the other substrates. Together, these data indicate that the competition experiment provides more detailed insight into complex formation than the separate experiments, revealing the preferentially formed ternary complex and thus best PROTAC substrate. The preference for Brd4^BD2^ observed by nMS fits to the increased half-life of the respective ternary complex (130s) as compared to that with Brd3^BD2^ (6s) and Brd4^BD1^ (<1s) (**Table 1** and ref. [12]). In fact, the lower half-life of the complex with Brd3^BD2^ is thought to be the reason for the lower degradation efficiency of Brd3 with respect to Brd4 in cells, despite similar binding affinity [12, 21, 22]. When the same MS experiments are performed with AT1, which has higher specificity for Brd4^BD2^, the signal intensity for the complex containing Brd4^BD2^ is higher than the other complexes, in both the low-competition and high-competition experiment (**Fig. 3c** and **Fig. 3d**). This reflects the preference of the formation of the VCB:AT1:Brd4^BD2^ complex over other substrate complexes.

**Figure 3.**
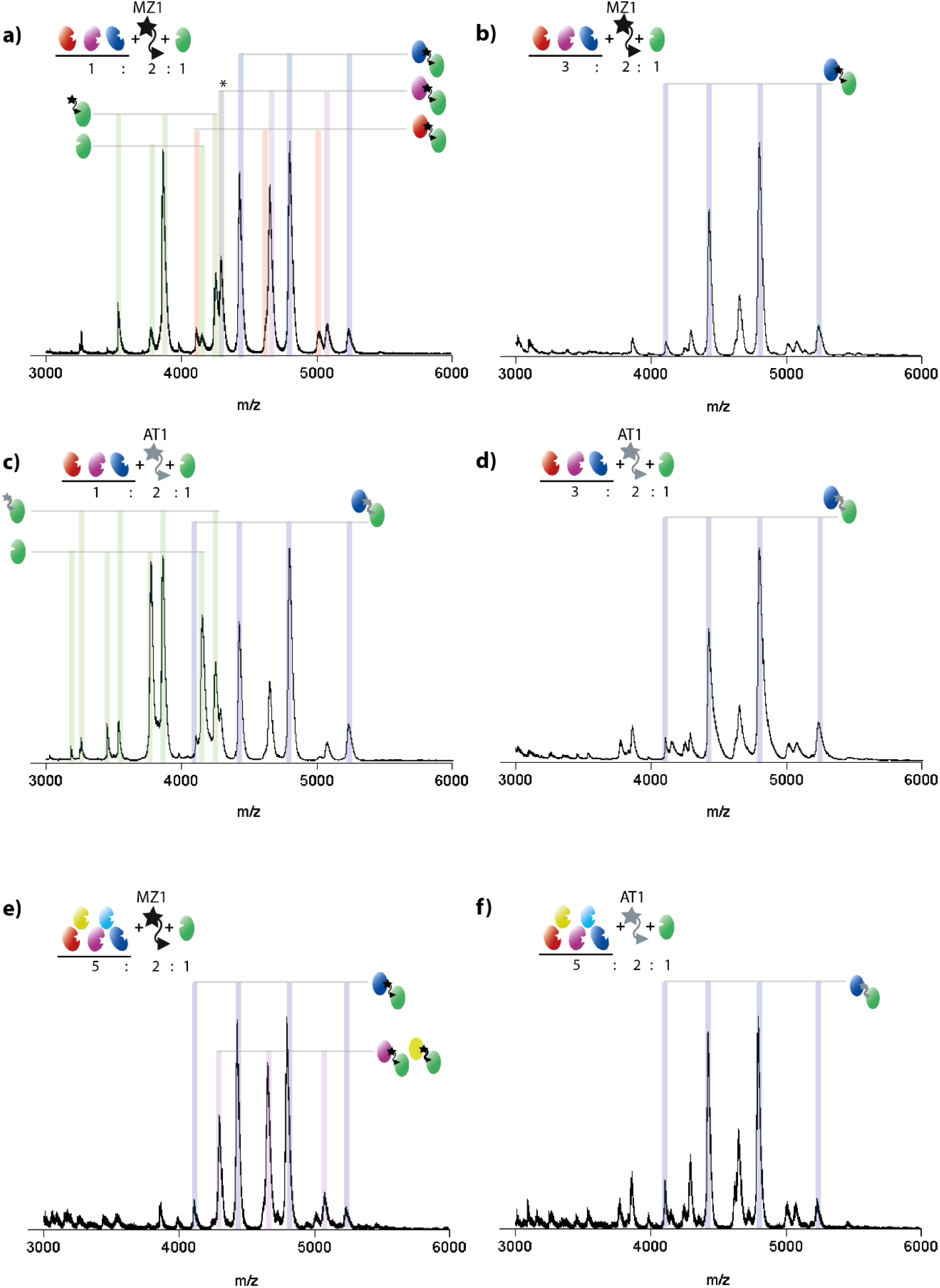
nESI spectra of VCB, PROTAC and a mixture of Bromodomain substrates. (a) VCB (5 μM), MZ1 (10 μM), equimolar mixture of Brd4^BD2^ (blue), Brd3^BD2^ (purple) and Brd4^BD1^ (red) (total Brd concentration 5 μM). (b) as a, but total Brd concentration 15 μM. (c, d) as a and b respectively, but PROTAC is AT1. (e) VCB (2.5uM), MZ1 (5uM), and a mixture of five Bromodomain substrates; Brd4^BD2^, Brd3^BD2^, Brd2^BD2^ (yellow), Brd4^BD1^, BrdT (cyan), total substrate concentration 12.5 μM. (f) as e, but PROTAC is AT1. Peaks corresponding to the most intense species are labelled, and fully annotated versions are given in **Fig S14.**

We next analyzed E3:P:S mixtures of even higher complexity, containing 5 bromodomain substrates and either MZ1 (**Fig. 3e**) or AT1 (**Fig. 3f**). Peaks can be separated for complexes containing Brd4^BD2^, Brd4^BD1^ and BrdT. The mass of Brd2^BD2^ is very close to that of Brd3^BD2^ (13 351 Da vs. 13 279 Da) and therefore complexes containing these proteins cannot be distinguished from one another. It is, however, clear from the spectra with both PROTACs that the complex containing Brd4^BD2^ has the highest intensity, inferring that this is the most favorable interaction. Additionally, the difference in intensity between Brd4^BD2^ and the next most intense peaks is bigger for the sample containing AT1 (**Fig. 3f**) with respect to MZ1 (**Fig 3e**), further demonstrating the higher specificity of this PROTAC. Such nMS experiments would be highly informative when screening proteins that are recruited by a certain PROTAC. Even if not every protein can be distinguished, as is the case for Brd3^BD2^ and Brd2^BD2^, the number of potential interactors can be greatly reduced for further investigation. Measuring the substrate proteins in mixtures is more time-effective than separate measurements and has the added advantage of providing information on competition between substrates forming the ternary complexes. Given the remarkable resolution of nMS, even small size differences in POIs, for instance introduced by adding short peptide tags, could be resolved, allowing the analysis of even more complex substrate sets as in the current analysis.

To conclude, we have demonstrated, for the first time, that nMS is an effective technique to investigate PROTAC-mediated protein complexes. We can determine differences in specificity of a PROTAC towards different proteins and can measure ternary complex formation of different substrates in a single experiment, which is highly beneficial in the generation of new PROTAC molecules. Whilst SPR and ITC remain the most appropriate methods for obtaining kinetic and thermodynamic data, we envision that nMS will become a popular tool in PROTAC development owing to its fast measurement time, straight-forward data analysis and ability to detect different species in equilibrium. Moreover, nMS bears the unique advantage to perform competition experiments, directly comparing potential substrates and various PROTACs to yield the most efficient degrading system.

## Online Methods

### Protein expression and purification

BRD2^BD2^ BRD3^BD2^, BRDt, BRD4^BD1^ and BRD4^BD2^ were expressed and purified as described by Filippakopoulos et al. [26] with final concentrations of 10.2 mg/mL (10 mM Hepes, 500 mM NaCl, 5% Glycerin, pH 7,5), 16 mg/mL (25 mM Hepes, 150 mM NaCl, 5 mM DTT, pH 7,5) 39.5 mg/mL (10 mM Hepes, 500 mM NaCl, 10 mM DTT, 5% Glycerin, pH 7,4), 13.4 mg/mL (50 mM Hepes, 500 mM NaCl, 5% Glycerin, pH 7,5) and 19 mg/mL (10 mM Hepes, 100 mM NaCl, 10 mM DTT, pH 7,5) respectively. Human VHL (54–213), ElonginC (17–112) and ElonginB (1–104) were co-expressed as described previously [5].All protein sequences are provided in **Table S2**.

### Sample preparation for native MS experiments

PROTACs were provided in a 10mM solution in DMSO, which was diluted 100× in water (100 μM, 1% DMSO). This was further diluted to 2× the working concentration using 1% DMSO in water, to ensure constant DMSO concentration across all experiments. Proteins were buffer exchanged into ammonium acetate using BioRad Micro Bio-Spin 6 Columns and the concentrations were measured with a Bradford Assay. Unless described otherwise, 20 μM substrate and 20 μM E3 ligase were mixed in an equimolar concentration (10 μM each) and added to an equivalent volume of PROTAC stock, to give final solution conditions of 5 μM substrate, 5 μM VCB, 5-10 μM PROTAC in 100 mM ammonium acetate, 0.5% DMSO.

### Mass spectrometry measurements

Native mass spectrometry experiments were carried out on a Synapt G2S*i* instrument (Waters, Manchester, UK) with a nanoelectrospray ionisation source. Mass calibration was performed by a separate infusion of NaI cluster ions. Solutions were ionised through a positive potential applied to metal-coated borosilicate capillaries (Thermo Scientific). The following instrument parameters were used for PROTAC complexes; capillary voltage 1.3 kV, sample cone voltage 80 V, extractor source offset 60 V, source temperature 40 °C, trap gas 3 mL/min. For individual proteins, the capillary voltage was set to 1.1 kV, sample cone voltage 40V, extractor source offset 30V, source temperature 40 °C and trap gas 2 mL/ min. Data were processed using Masslynx V4.1 and GraphPad Prism 8.1.1. To determine the estimated ratio of signal corresponding to each species, the relative intensity of peaks involved in the comparison were summed, and the sum of peaks for a particular species was divided by the sum of the total peaks.

## Acknowledgements

We are grateful to Karl Mechtler and the Vienna Biocenter Core Facilities for providing the mass spectrometry infrastructure required to perform this research. We are also very appreciative of Dr Sophie Rebecca Harvey and Dr Lukasz Migas for critical reading of the manuscript and their useful comments. RB acknowledges the Austrian Science Fund for the receipt of a Lise Meitner Postdoctoral Fellowship (project number M2334). The IMP is funded by Boehringer Ingelheim.

## Author Contributions

R.B. and T.C. designed the experiments, which were performed by R.B.. Data analyses were performed by R.B, D.R., K.R., P.E., A.M. and T.C.. R.B. and T.C. drafted and edited the manuscript.

