## supplementary information for "Native mass spectrometry can effectively predict PROTAC efficacy"

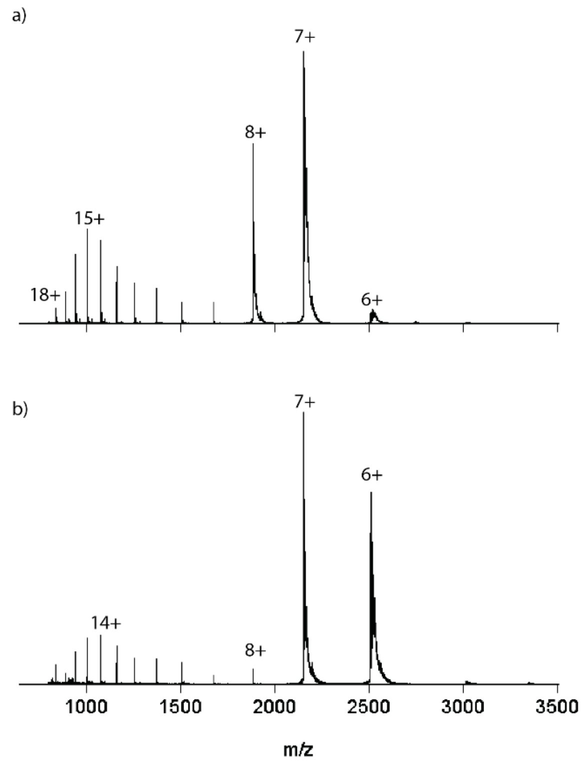


**Supplementary Information Figure 1.** nESI-MS of Brd4$^{BD2}$ sprayed from (a) 100mM ammonium acetate and (b) 100mM ammonium acetate, 0.5% DMSO. In (a), Brd4$^{BD2}$presents in charge states $[M+6H]^{6}$ to $[M+18H]^{18+}$, with most of the intensity in charge states $[M+7H]^{7+}$ and $[M+8H]^{8+}$. Such a distribution of charge states corresponds to a protein that exists in both extended and compact conformations in solution; the low charge states correspond to a folded conformation that has a limited number of solvent-accessible protonatable sites, whilst the higher charge states correspond to more extended conformations. The addition of 0.5% DMSO, which was added to account for the solvent present in the PROTAC sample, shifts the protein to lower charge states, reasons for which have been discussed by Chan et al. [1].

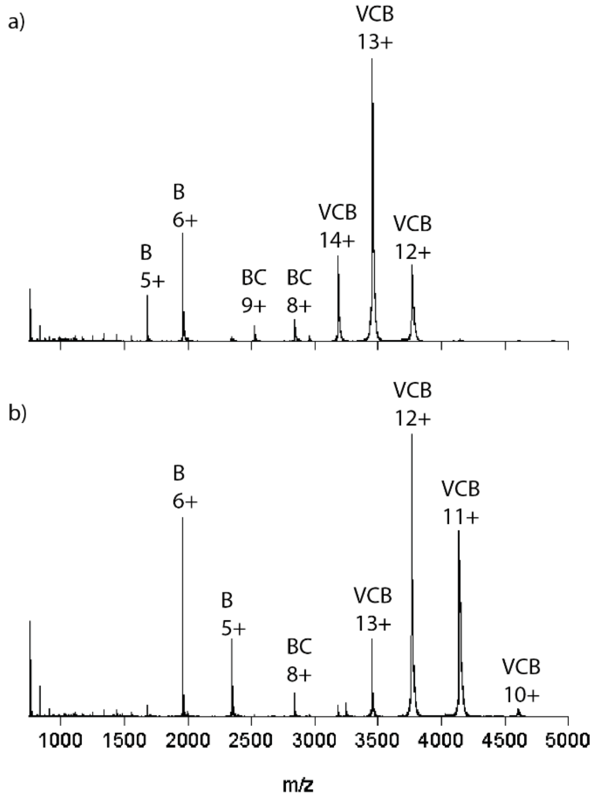


**Supplementary Information Figure 2.** nESI-MS of VCB sprayed from (a) 100mM ammonium acetate and (b) 100mM ammonium acetate, 0.5% DMSO. In (a), the trimeric VCB complex presents in charge states $[M+12H]^{12+}$ to $[M+14H]^{14+}$. Elongin B- elongin C dimer (labelled BC) is present at low relative intensity in charge states $[M+8H]^{8+}$ and $[M+9H]^{9+}$, and monomeric elongin B (labelled B) is present in charge states $[M+5H]^{5+}$ and $[M+6H]^{6+}$. When 0.5% DMSO is present in the solution, the same species are present, at lower charge states.

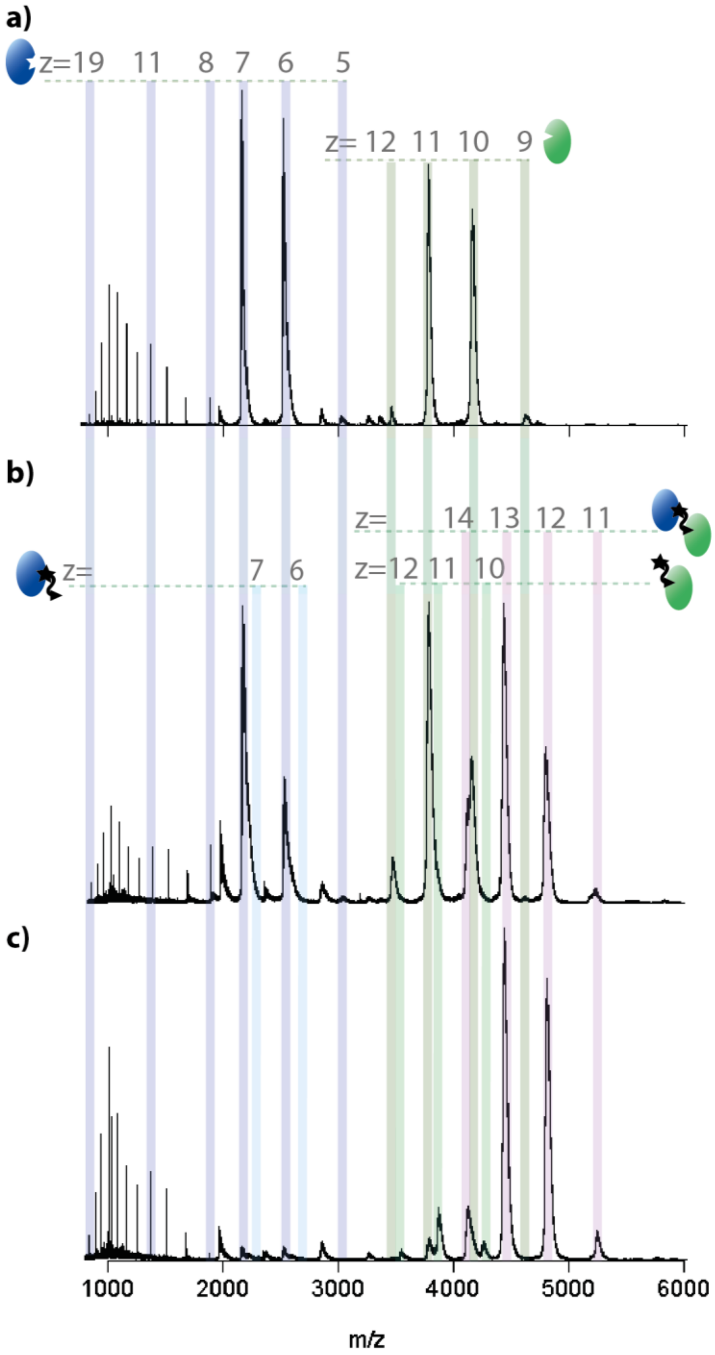


**Supplementary Information Figure 3.** nESI-MS of Brd4$^{BD2}$ (5 µM) and VCB (5 µM) sprayed from ammonium acetate (100 mM, pH 6.8) and 0.5% DMSO at MZ1 (1002 Da) concentrations of 0 µM (a), 5 µM (b) and 10 µM (c). Insets show estimated relative signal intensity of integrated peaks corresponding to *apo*-Brd4$^{BD2}$, binary MZ1-Brd4$^{BD2}$ complex, *apo*-VCB, binary MZ1-VCB complex and ternary complex Brd4$^{BD2}$-MZ1-VCB.

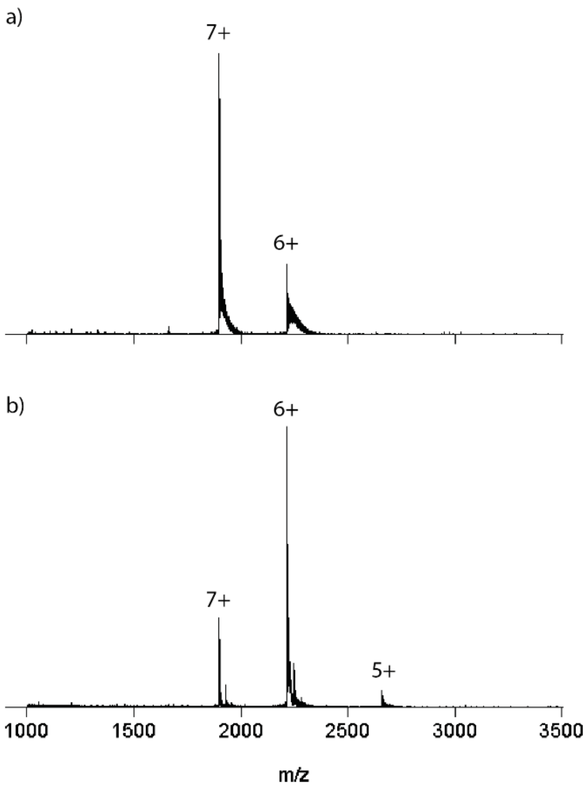


**Supplementary Information Figure 4.** nESI-MS of Brd3$^{BD2}$ sprayed from (a) 100mM ammonium acetate and (b) 100mM ammonium acetate, 0.5% DMSO. In (a), Brd3$^{BD2}$ presents in just two charge states; $[M+6H]^{6+}$ and $[M+7H]^{7+}$. This indicates that Brd3$^{BD2}$ exists in a more compact conformation than Brd4$^{BD2}$. Again, the addition of DMSO results in lower charge states.

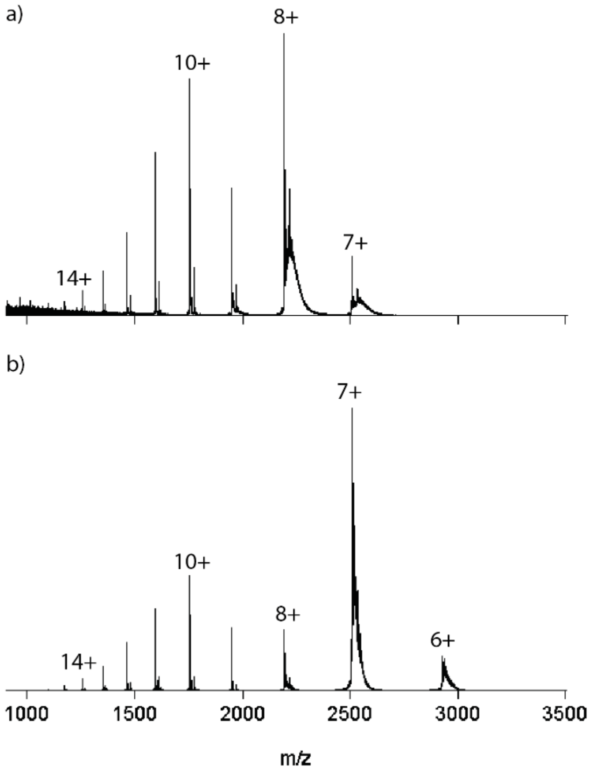


**Supplementary Information Figure 5.** nESI-MS of Brd4$^{BD1}$ sprayed from (a) 100mM ammonium acetate and (b) 100mM ammonium acetate, 0.5% DMSO. In (a), Brd4$^{BD1}$presents in charge states $[M+7H]^{7+}$ to $[M+14H]^{14+}$, with a more even distribution of intensity in the lower and higher charge states than Brd4$^{BD2}$, corresponding to a more disordered protein. (b) 0.5% DMSO causes a shift towards lower charge states.

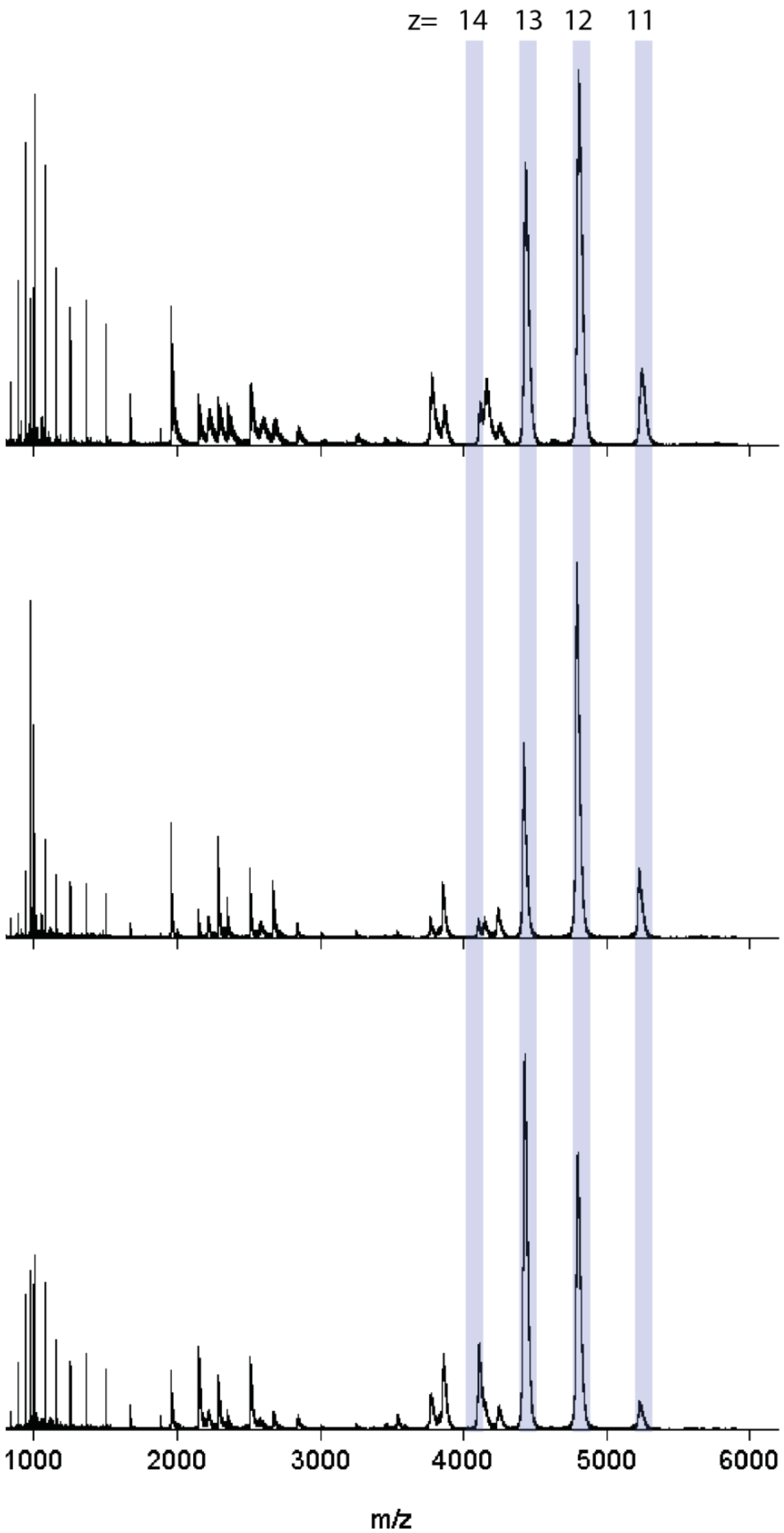


**Supplementary Information Figure 6**. Three technical nESI measurements of VCB (5µM) + $Brd4^{BD2}$ (5µM) + AT1 (10µM). Peaks corresponding to the ternary complex are highlighted.

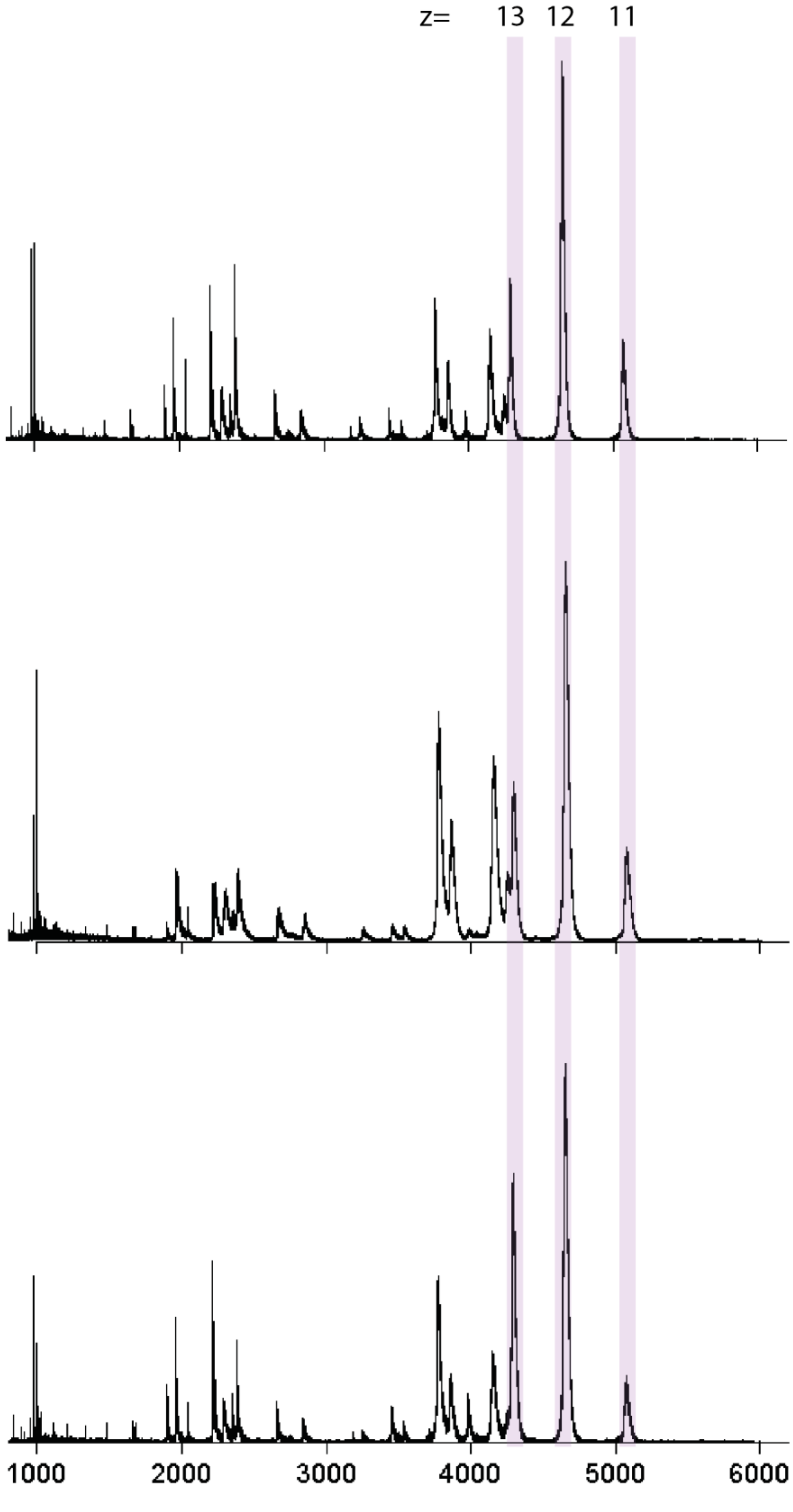


**Supplementary Information Figure 7**. Three technical nESI measurements of VCB (5µM) + Brd3$^{BD2}$ (5µM) + AT1 (10µM). Peaks corresponding to the ternary complex are highlighted.

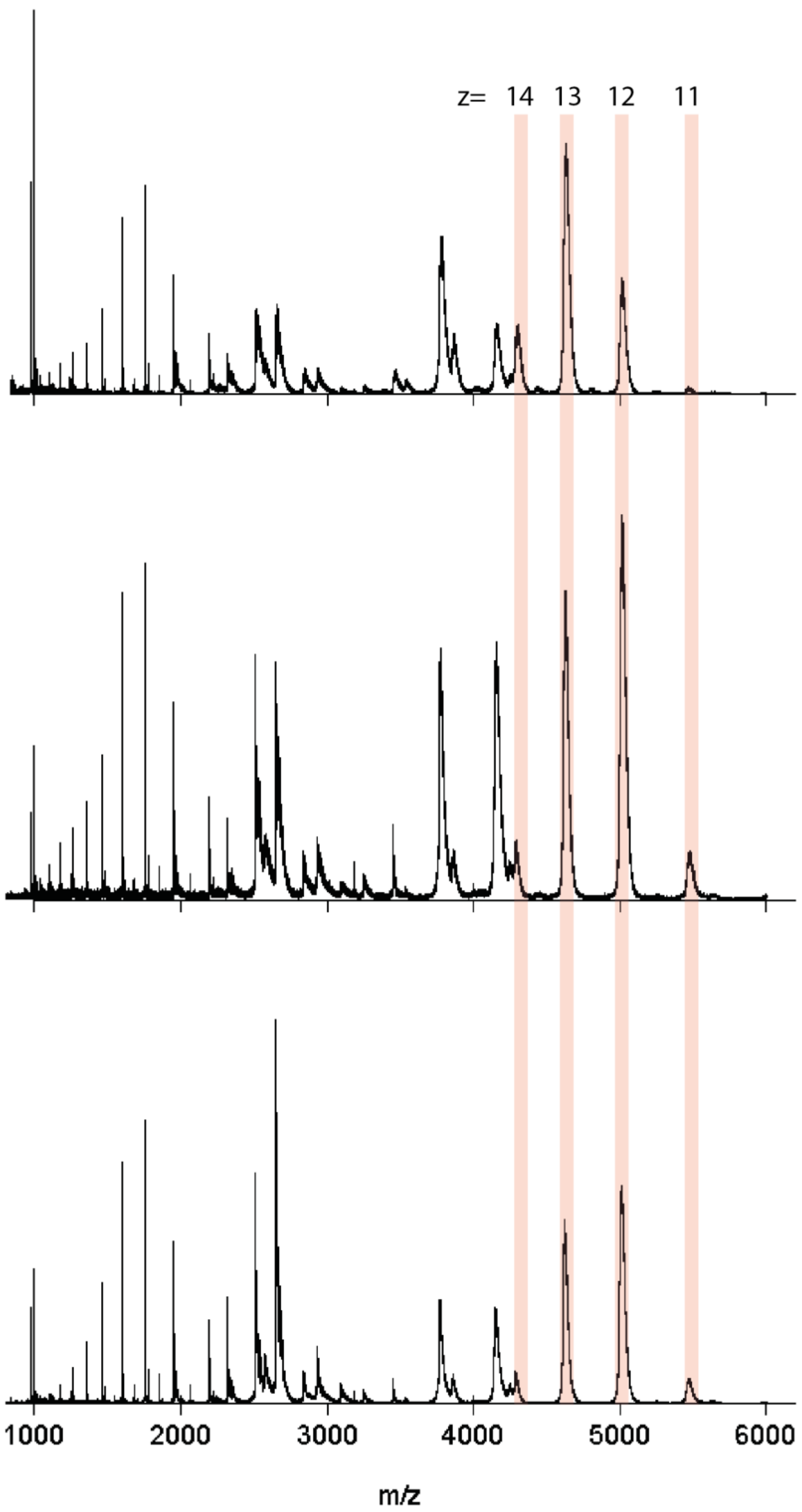


**Supplementary Information Figure 8**. Three technical nESI measurements of VCB (5µM) + Brd4[BD1] (5µM) + AT1 (10µM). Peaks corresponding to the ternary complex are highlighted.

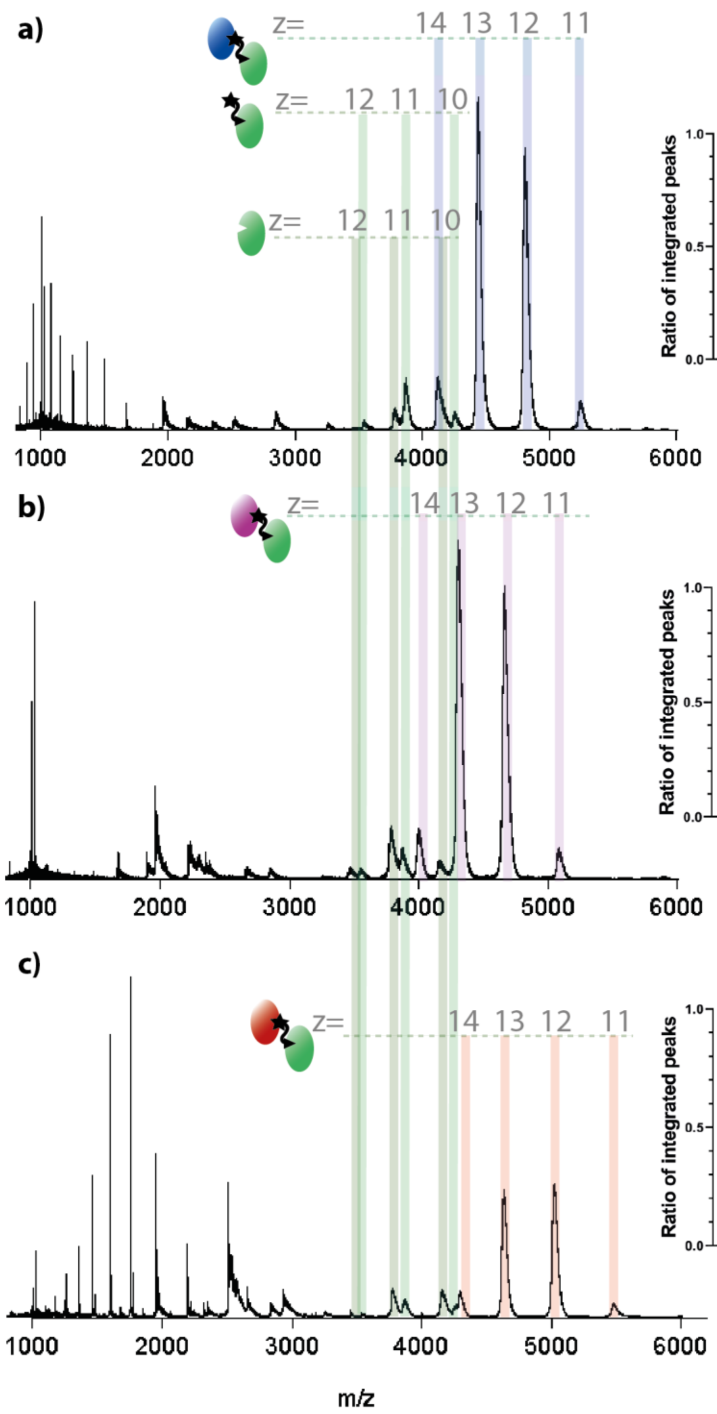


**Supplementary information Figure 9** nESI-MS of VCB (5 µM), MZ1 (10 µM) and Brd4$^{BD2}$ (5 µM, a), Brd3$^{BD2}$ (5 µM, b) or Brd4$^{BD1}$ (5 µM, c). Proteins are sprayed from a starting solution of ammonium acetate (100 mM, pH 6.8) and 0.5% DMSO. Inset; estimated relative intensity of summed peaks corresponding to *apo*- VCB, binary MZ1-VCB complex and ternary complex substrate-MZ1-VCB. Samples were analysed in triplicate and the error bars represent standard deviation of the relative peak intensity.

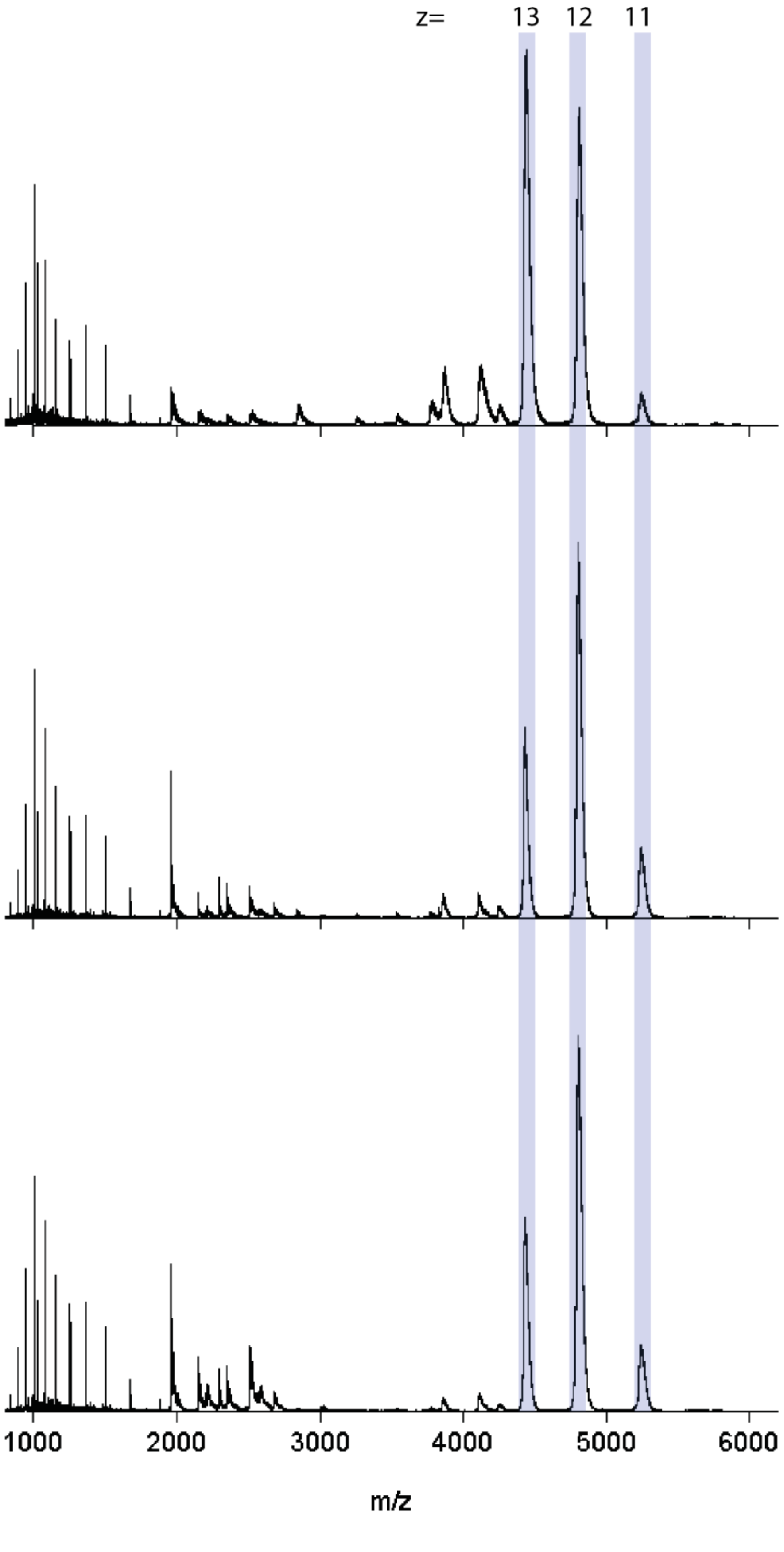


**Supplementary Information Figure 10**. Three technical nESI measurements of VCB (5µM) + Brd4$^{BD2}$ (5µM) + MZ1 (10µM). Peaks corresponding to the ternary complex are highlighted.

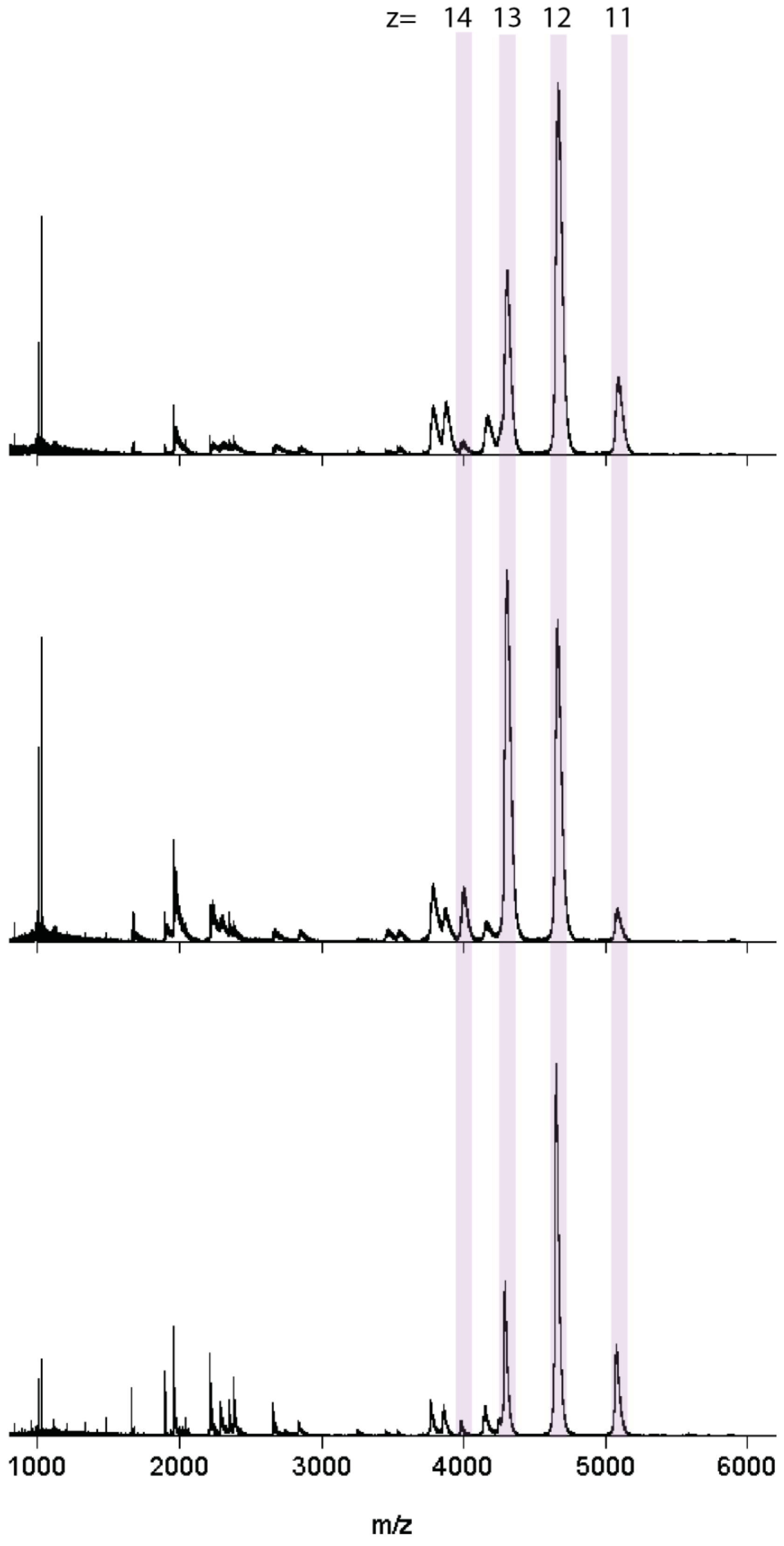


**Supplementary Information Figure 11**. Three technical nESI measurements of VCB (5µM) + Brd3$^{BD2}$ (5µM) + MZ1 (10µM). Peaks corresponding to the ternary complex are highlighted.

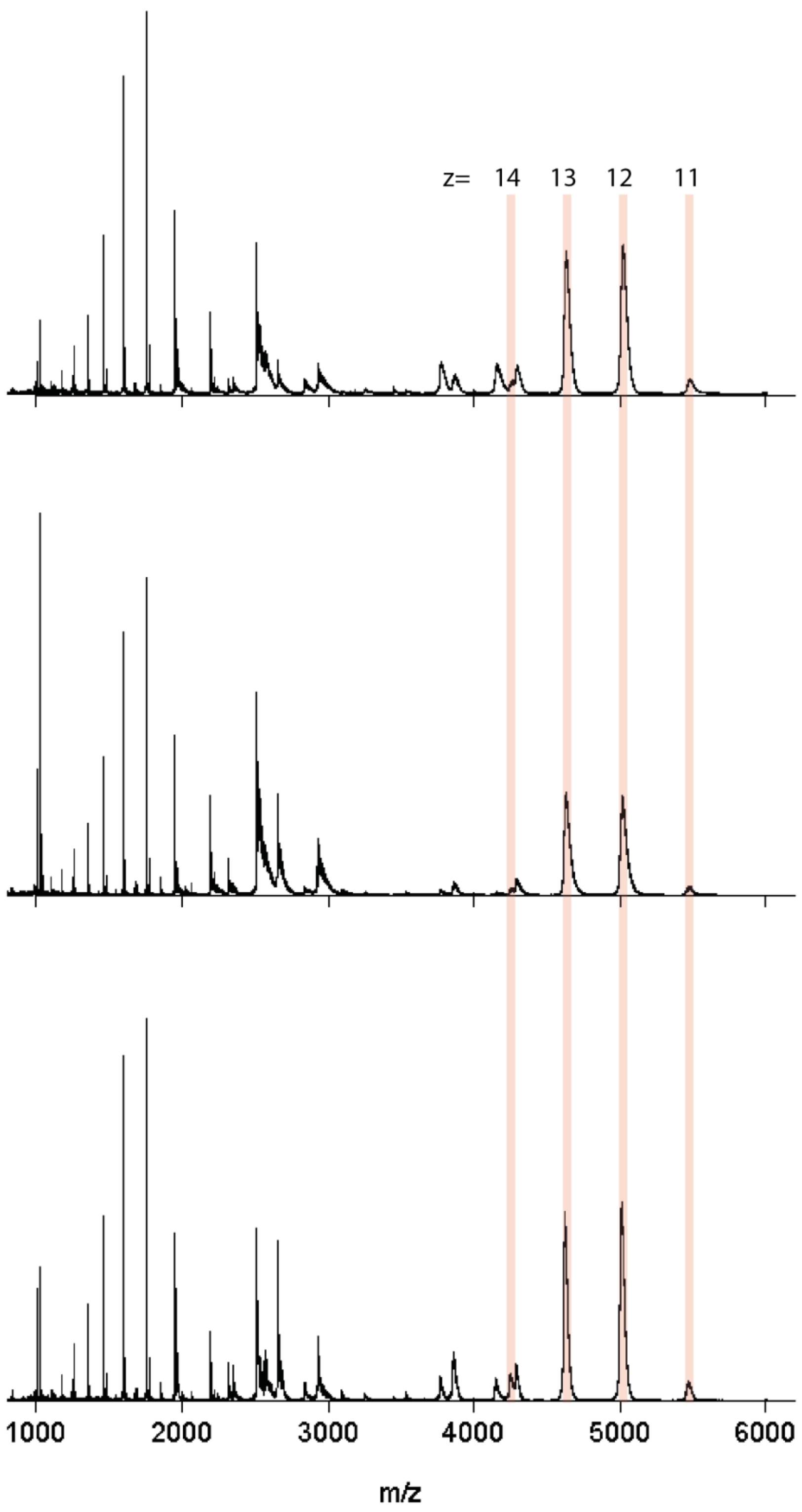


**Supplementary Information Figure 12**. Three technical nESI measurements of VCB (5µM) + Brd4$^{BD1}$ (5µM) + MZ1 (10µM). Peaks corresponding to the ternary complex are highlighted.

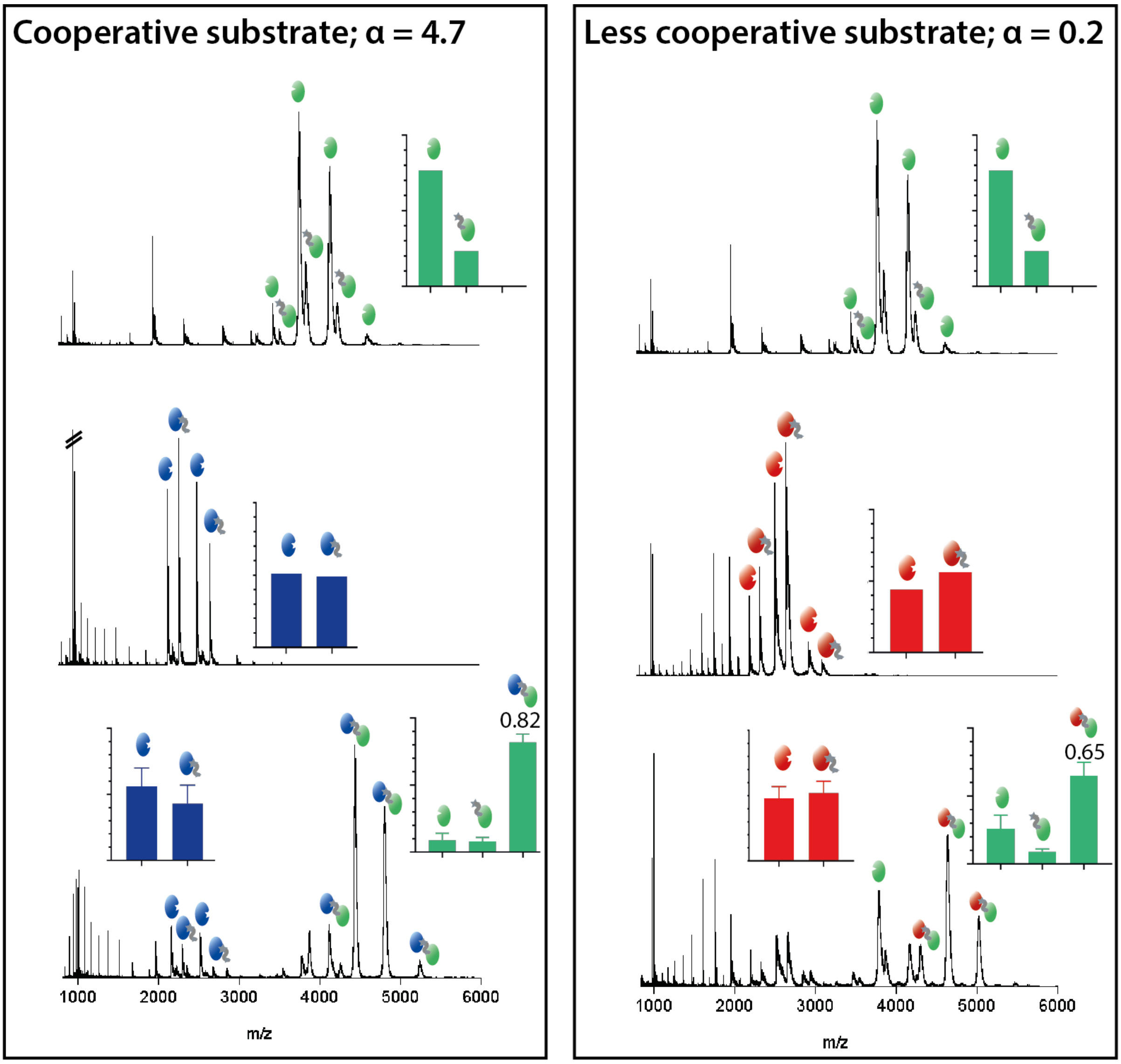


**Supplementary Information Figure 13:** Methods were also developed for testing PROTAC cooperativity.

Top; VCB (5 µM) + AT1 (10 µM). Middle; substrate protein (5 µM) + AT1 (10 µM). Bottom; VCB (5 µM) + AT1 (10 µM) + substrate protein (5 µM). Left; substrate is Brd4$^{BD2}$. Right; substrate protein is Brd4$^{BD1}$. Inset: estimated fraction of integrated peaks corresponding to the labelled species. For Brd4$^{BD2}$ and Brd4$^{BD1}$, only the peaks which correspond to $[M+6H]^{6+}$ and $[M+7H]^{7+}$ for Brd4$^{BD2}$ and $[M+5H]^{5+}$ $[M+6H]^{6+}$ and $[M+7H]^{7+}$ for Brd4$^{BD1}$ are used for the quantification. Insets in the top and middle panels correspond to single measurements, whilst those in the bottom panel are the average of three measurements, with error bars representing standard deviations.

In a cooperative PROTAC system, the ternary complex will form more readily than either of the binary complexes. We measured the two-component mixtures [5µM VCB + 10µM PROTAC] and [5µM substrate + 10µM PROTAC], followed by the three-component mixture [5µM VCB + 5µM substrate + 10uM PROTAC]. In a cooperative system, the formation of protein-PROTAC binary complex will be low with each protein, but the ternary complex formation will be high when all the components are present. This figure shows how native MS can determine cooperativity; binary complex formation between VCB and AT1 is low, below 20%, and the binary complex formation between $Brd4^{BD2}$ and AT1 and between $Brd4^{BD1}$ and AT1 is roughly 50% in both cases. When the three components are mixed together, the ternary complex is formed to a much higher extent with $Brd4^{BD2}$ than $Brd4^{BD1}$, due to cooperativity of this system [2].

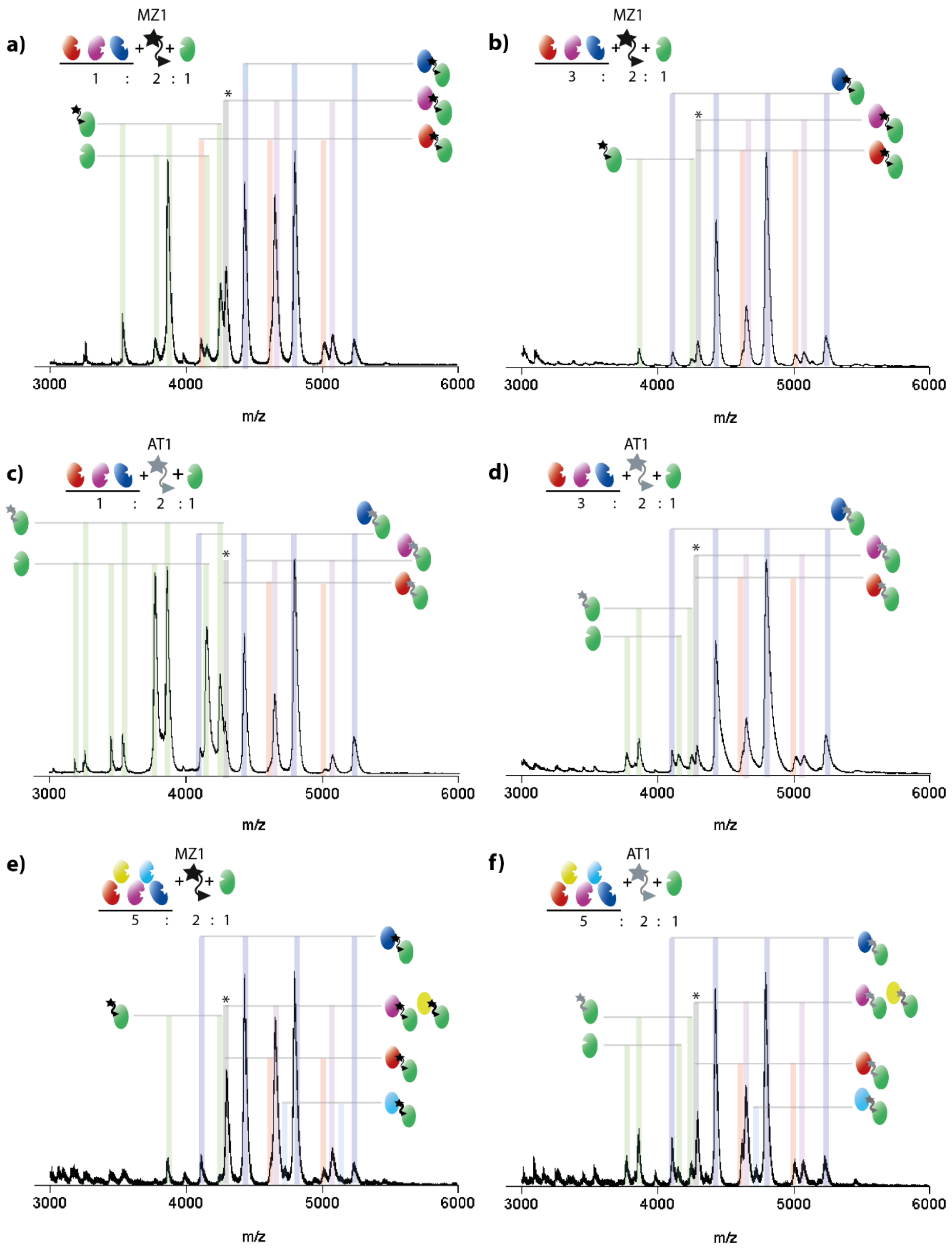


**Supplementary Information Figure 14** Fully annotated versions of nESI spectra shown in Figure 3 of VCB, PROTAC and a mixture of Bromodomain substrates. (a) VCB (5 µM), MZ1 (10 µM), equimolar mixture of Brd4$^{BD2}$ (blue), Brd3$^{BD2}$ (purple) and Brd4$^{BD1}$ (red) (total concentration 5 µM). (b) as a, but total Brd concentration 15 µM. (c, d) as a and b respectively, but PROTAC is AT1. (e) VCB (2.5uM), MZ1 (5uM), and a mixture of five Bromodomain substrates; Brd4$^{BD2}$,

Brd3$^{BD2}$, Brd2$^{BD2}$ (yellow), Brd4$^{BD1}$, BrdT (cyan), total substrate concentration 12.5 µM. (f) as e, but PROTAC is AT1.

| Species | Expected mass/ Da | Measured mass/ Da |
|---|---|---|
| **Brd4$^{BD2}$** | 15 036 (expected from sequence) | 15 036 |
| **Brd4$^{BD2}$ + AT1** | 16 007 | 16 007 |
| **VCB** | 41 373 (expected from sequence) | 41 376 |
| **VCB + AT1** | 42 348 | 42 528 ± 12$^{\$}$ |
| **Brd4$^{BD2}$ + AT1+ VCB** | 57 384 | 57 636.5 ± 33.5$^{\$}$ |

**Supplementary Information Table 1.** Expected and measured mass (taken from the apex of the peaks) of species present in Figure 1.

$^{\$}$ Increased measured mass c.f. expected mass is due to unresolved salt and solvent adducts bound to the protein. Importantly, for all species, the expected mass corresponds to the left-hand side of the m/z peaks (that have no retention of salt).

| Protein | Sequence |
|---|---|
| Brd4$^{BD1}$ | mhhhhhhssgvdlgtenlyfqsmNPPPPETSNPNKPKRQTNQLQYLLRVVLKTLWKHQFAWPF QQPVDAVKLNLPDYYKIIKTPMDMGTIKKRLENNYYWNAQECIQDFNTMFTNCYIYNKPG DDIVLMAEALEKLFLQKINELPTEE |
| Brd4$^{BD2}$ | smKDVPDSQQHPAPEKSSKVSEQLKCCSGILKEMFAKKHAAYAWPFYKPVDVEALGLHDY CDIIKHPMDMSTIKSKLEAREYRDAQEFGADVRLMFSNCYKYNPPDHEVVAMARKLQDVF EMRFAKMPDE |
| Brd3$^{BD2}$ | smKLSEHLRYCDSILREMLSKKHAAYAWPFYKPVDAEALELHDYHDIIKHPMDLSTVKRKM DGREYPDAQGFAADVRLMFSNCYKYNPPDHEVVAMARKLQDVFEMRFAKMP |
| VCB; VHL subunit | gsMEAGRPRPVLRSVNSREPSQVIFCNRSPRVVLPVWLNFDGEPQPYPTLPPGTGRRIHSYR GHLWLFRDAGTHDGLLVNQTELFVPSLNVDGQPIFANITLPVYTLKERCLQVVRSLVKPENY RRLDIVRSLYEDLEDHPNVQKDLERLTQERIAHQRMGD |
| VCB; Elongin B subunit | MDVFLMIRRHKTTIFTDAKESSTVFELKRIVEGILKRPPDEQRLYKDDQLLDDGKTLGECGFT SQTARPQAPATVGLAFRADDTFEALCIEPFSSPPELPDVMK |
| VCB; Elongin C subunit | MMYVKLISSDGHEFIVKREHALTSGTIKAMLSGPGQFAENETNEVNFREIPSHVLSKVCMYF TYKVRYTNSSTEIPEFPIAPEIALELLMAANFLDC |
| Brd2 | smGKLSEQLKHCNGILKELLSKKHAAYAWPFYKPVDASALGLHDYHDIIKHPMDLSTVKRK MENRDYRDAQEFAADVRLMFSNCYKYNPPDHDVVAMARKLQDVFEFRYAKMPD |
| BrdT | smNTKKNGRLTNQLQYLQKVVLKDLWKHSFSWPFQRPVDAVKLQLPDYYTIIKNPMDLNTI KKRLENKYYAKASECIEDFNTMFSNCYLYNKPGDDIVLMAQALEKLFMQKLSQMPQEE |

**Supplementary Information Table 2** sequences of all proteins used in the study


1. Chan, D.S.-H., et al., *Insight into Protein Conformation and Subcharging by DMSO from Native Ion Mobility Mass Spectrometry.* ChemistrySelect, 2016. **1**(18): p. 5686-5690.
2. Gadd, M.S., et al., *Structural basis of PROTAC cooperative recognition for selective protein degradation.* Nature Chemical Biology, 2017. **13**: p. 514.
